# Complement Factor H (Y402H) polymorphism for age-related macular degeneration alters retinal lipids

**DOI:** 10.64898/2026.01.06.697989

**Authors:** Peng Shang, Johnson Hoang, Elise Hong, Zhaohui Geng, Helena Ambrosino, Eric Abnoosian, Xiaoyu Zhu, Min Ma, Nova A. Wei-Navarro, Beau Webber, Jun Qu, Sandra R. Montezuma, Martin-Paul Agbaga, James R Dutton, Deborah A. Ferrington

## Abstract

Age-related macular degeneration (AMD), the leading cause of blindness in the elderly, is associated with multiple risk factors and involves death of the retinal pigment epithelium (RPE). We investigated how the Y402H polymorphism of Complement Factor H (CFH) and cigarette smoke extract (CSE), major AMD genetic and environmental risks, affect lipid metabolism in RPE differentiated from induced pluripotent stem cells (iPSC-RPE) that were derived from human donors genotyped for low-risk (LR) or high-risk (HR) CFH. Results from discovery-based (lipidomics, proteomics) and targeted (mitochondrial fatty acid oxidation (FAO)) assays found significant genotype-dependent differences under basal conditions include higher free fatty acids and cholesterol esters in HR cells. CSE induced differences in proteins regulating lipid handling, lipolysis, and inflammation. Lower FAO in HR cells was observed in multiple donors and pairs of parent/isogenic edited lines compared with LR lines. CSE induced lipid accumulation, lipid composition remodeling, and upregulation of proteins involved in lipid synthesis/hydrolysis, production of bioactive lipid mediators, and metabolism of ceramide and cholesterol. These results elucidate putative mechanisms driving pathology in RPE harboring CFH Y402H.

## Introduction

Age-related macular degeneration (AMD), the leading cause of vision loss in the elderly, affects ∼25% of individuals age 75 years and increases in prevalence to 40% in the following decade of life (1). AMD manifests as a loss of central vision, which results in difficulties with essential daily functions, such as reading, driving, and face recognition. The loss of central vision is due to the dysfunction and death of the retinal pigment epithelium (RPE) and subsequent loss of the light-sensing photoreceptors in the macula. The RPE is a monolayer of cells located between the retinal photoreceptors and the choroid, which contains blood vessels supplying the outer retina with oxygen and nutrients (2). The RPE performs multiple functions required to maintain vision, including transport of nutrients and oxygen from the choroid to the photoreceptors, secretion of molecules that are crucial for the health and integrity of the surrounding tissue, and the daily phagocytosis of the oldest portion of the photoreceptor outer segment membranes, thereby eliminating damaged segments in these light-sensing cells (2).

Multiple risk factors have been associated with AMD development and progression, including age, genetic background, lifestyle and environment, especially smoking. The significant contribution of genetics to AMD risk was underscored in a large population-based study that reported genetics account for approximately 70% of the risk for developing advanced AMD (3). This genome-wide association study (GWAS) of AMD patients identified 52 common and rare single nucleotide polymorphisms (SNP) in 34 high-risk loci involving different biological pathways, including complement, lipid transport and processing, extracellular matrix remodeling, and angiogenesis (3). However, the biological consequences of the polymorphisms in these different biological processes and how they contribute to AMD pathology are still not fully elucidated.

The goal of the current study is to define the interplay between two biologically distinct pathways linked to AMD: lipid metabolism and the complement system. Evidence supporting dysregulation of retinal lipid homeostasis as a key contributor to AMD pathology include identification of SNPs in genes regulating cholesterol efflux (ABCA1, ABCG1), lipid transport (APOE), and metabolism (LIPC, CETP, LPL) (4). Direct evidence of disrupted lipid handling in AMD is indicated by the presence of lipid/cholesterol-rich drusen located between the RPE and Bruch’s membrane (5) and subretinal deposits between the RPE and photoreceptors (6), which are early lesions associated with AMD and significant risk factors for developing advanced AMD (7). Retinal lipid handling is a tightly regulated process, in part due to the involvement of lipids in essential cellular functions. As the major structural component of cell and organelle membranes, the lipid bilayer is composed of multiple lipid classes (phospholipids, and sterols, mainly cholesterol) that impart biophysical properties uniquely required for cellular membrane and organellar function (8). Lipids also serve as secondary messengers by producing bioactive lipids that regulate signaling pathways, such as inflammation, calcium release, and cell death. They are also an important source of energy, as free fatty acids are the substrates for fatty acid oxidation in the mitochondria through β-oxidation (9, 10). Each of these processes is precisely controlled by specific enzymes that respond to external or internal signals that arise due to changes in the cellular environment. Defects or imbalance in any of these lipid-based processes could lead to RPE dysfunction and eventual cell death, which are attributes of AMD.

Polymorphisms in at least seven genes of the complement pathway have been associated with AMD, including complement factor H (CFH), a negative regulator of the alternative complement pathway, which promotes clearance of cellular debris and kills invading pathogens (11). The CFH Y402H SNP (rs1061170), found in ∼50% of AMD patients, has been shown to increase AMD risk 2.7- to 7.4-fold in individuals possessing one or two risk alleles (12–14). The change of a single nucleotide in the CFH gene (T to C) results in substitution of tyrosine for histidine at position 402 in the CFH protein. Overactivation of the complement cascade due to the reduced ability of the Y402H CFH protein to inhibit complement activation can lead to chronic inflammation and lysis of host cells via the formation of the membrane attack complex on cell outer membranes (15). Chronic inflammation and retinal cell death are well-known characteristics of AMD pathology and could result from extracellular CFH dysregulation.

While the canonical role for extracellular complement, as described above, has been well-studied, recent studies provide evidence of an emerging role for intracellular complement that includes regulation of multiple cellular pathways. Investigations from our laboratory using RPE differentiated from induced pluripotent stem cells (iPSC-RPE) reported cells with the high risk CFH Y402H genotype had decreased mitochondrial function and showed significant differences in their response to oxidative stress compared with iPSC-RPE harboring the low-risk CFH genotype (16, 17). Other laboratories also have reported distinct genotype-specific cellular responses in iPSC-RPE with this high-risk CFH SNP, including lysosomal damage and reduced autophagy, elevated inflammation, and of relevance to the current investigation, lipid deposition (18, 19).

In this study, we continue using iPSC-RPE cell lines derived from human donors genotyped for the CFH SNP (rs1061170) to investigate how the high risk CFH genotype affects lipid homeostasis in RPE. We include exposure to cigarette smoke extract (CSE) as a relevant model of environmental stress. Our prior study showed CSE induced significant oxidative stress, altered the metabolic and protein profiles, and reduced mitochondrial function in iPSC-RPE (17). Smoking tobacco, a lifestyle choice and primary environmental risk factor associated with AMD, increases the risk for AMD 3- to 4-fold in past and current smokers, respectively (20). In smokers homozygous for the CFH Y402H high risk allele, the odds ratio increases to 34-fold for advancing to late-stage AMD compared with non-smokers without this CFH risk SNP (21).

These results show the significant multiplicative effect of combining prominent genetic and environmental risk factors for AMD. Layering AMD risk factors allowed us to tease out genotype-dependent differences in lipid homeostasis in the untreated basal state, as well as the response to chronic CSE-induced stress. These analyses inform our understanding of how genetic and environmental factors alter lipid homeostasis and cause the increased risk for AMD in individuals carrying the CFH risk genotype.

## Results

### CFH Y402H SNP and CSE treatment significantly alter RPE protein and lipid profiles

This study used iPSC-RPE cell lines derived from human donors genotyped for the CFH Y402H SNP associated with high risk for AMD (rs1061170) and evaluated for AMD presence and severity using the Minnesota Grading System (MGS) (22). Donor demographics, genotype, and AMD stage are provided in Table 1. Each experimental group consisted of iPSC-RPE differentiated from iPSC lines derived from individual donors with either the low-risk genotype (LR group) or the high-risk genotype (HR group). Note that all donors were homozygous for their respective CFH Y402H SNP (LR, TT or HR, CC). Cell lines used for different analyses (proteomics, lipidomics, fatty acid oxidation) and the donor’s average age for each group are indicated (Table 1). Average donor age (also provided for each assay) for groups in each experiment was not significantly different.

**Table 1.** Donor demographics and analysis for iPSC-RPE.

| **iPSC-RPE Line**<br>**ID** | **Proteomics** | **Lipidomics** | **Fatty Acid**<br>**Oxidation** | **Age<sup>a</sup>/Sex<sup>b</sup>** | **MGS<sup>c</sup>** | **CFH**<br>**Genotype<sup>d</sup>** |
| --- | --- | --- | --- | --- | --- | --- |
| MGS1-1418 | ✓ | ✓ | ✓ | 54/M | MGS1 | TT |
| MGS1-1351 |  | ✓ |  | 67/M | MGS1 | TT |
| MGS1-0237 | ✓ | ✓ |  | 77/F | MGS1 | TT |
| MGS2-2275 | ✓ | ✓ | ✓ | 68/M | MGS2 | TT |
| MGS2-0179 | ✓ | ✓ | ✓ | 71/F | MGS2 | TT |
| MGS2-1389 | ✓ | ✓ | ✓ | 75/M | MGS2 | TT |
| MGS2-1935 | ✓ | ✓ |  | 80/F | MGS2 | TT |
| MGS2-0833 | ✓ | ✓ | ✓ | 83/F | MGS2 | TT |
| MGS3-0277 | ✓ | ✓ | ✓ | 80/F | MGS3 | TT |
| MGS3-0011 |  | ✓ |  | 83/F | MGS3 | TT |
| MGS1-1974 | ✓ | ✓ | ✓ | 62/F | MGS1 | CC |
| MGS1-0987 | ✓ | ✓ | ✓ | 82/F | MGS1 | CC |
| MGS2-0984 | ✓ | ✓ | ✓ | 67/F | MGS2 | CC |
| MGS2-1825 | ✓ | ✓ |  | 70/F | MGS2 | CC |
| MGS2-0024 | ✓ | ✓ | ✓ | 75/F | MGS2 | CC |
| MGS3-0698 | ✓ | ✓ | ✓ | 72/F | MGS3 | CC |
| MGS3-0276 | ✓ | ✓ | ✓ | 75/F | MGS3 | CC |
| MGS3-0503 | ✓ | ✓ |  | 83/F | MGS3 | CC |
| MGS3-1404 | ✓ |  |  | 84/M | MGS3 | CC |

|  | **Proteomics** | **Lipidomics** | **Fatty Acid Oxidation** |
| --- | --- | --- | --- |
| **TT** | (73.5 ± 8.7), 5F/3M | (73.8 ± 8.6), 6F/4M | (71.8 ± 9.4), 3F/4M |
| **CC** | (74.4 ± 7.1), 8F/1M | (73.3 ± 6.7), 8F/0M | (72.1 ± 6.4), 6F/0M |

| **Parent and isogenic iPSC-RPE for fatty acid oxidation** | **Age<sup>a</sup>/Sex<sup>b</sup>** | **MGS<sup>c</sup>** | **CFH Genotype<sup>d</sup>** |
| --- | --- | --- | --- |
| MGS1-0237- parent | 77/F | MGS1 | TT |
| MGS1-0237-edited | 77/F | MGS1 | CC |
| MGS2-0984-parent | 67/F | MGS2 | CC |
| MGS2-0984-edited | 67/F | MGS2 | TT |

Age (Mean ± SD); Sex (F: Female; M: Male)
<sup>a</sup> Age of donor (years) used to generate iPSC-RPE.
<sup>b</sup> Sex of donor, F = female, M = male.
<sup>c</sup> Minnesota Grading System (MGS) was used to evaluate the stage of AMD in eye bank eyes (22). No AMD = MGS1; MGS2 (early AMD) and MGS3 (intermediate AMD).
<sup>d</sup> Complement Factor H (CFH) genotype for rs106117: low risk = TT, high risk = CC.

To model an environment of chronic oxidative stress, iPSC-RPE cells from multiple CFH LR and HR donors were exposed to cigarette smoke extract (CSE) for two weeks. At the end of CSE exposure, both treated and vehicle (DMSO) control cells were collected for either shotgun lipidomics (LR=10, HR=8 lines from individual donors) or proteomics analyses (LR=7, HR=8 lines from individual donors). The goal of the combined “omics” approach is to obtain an unbiased overview of how genotype and CSE treatment affect the lipid and protein profile in the RPE. The proteomics data was used to complement the lipid analysis by providing insight into the molecular mechanisms that drive changes in RPE lipid metabolism. Lipidomics analysis identified and quantified 1,170 lipid species, comprising 17 lipid subclasses categorized by their chemical backbone and esterified or amide-linked fatty acids. The proteomics analysis identified and quantified 9432 proteins across all experimental groups. Repeated-measure (RM) two-way ANOVA was performed on the two datasets to determine the effects of the CFH Y402H SNP, CSE treatment, and the interaction of these two factors on each lipid species and protein.

Statistical analysis of each dataset revealed significant effects for both CFH genotype and CSE exposure, reflecting underlying differences between CFH HR and LR RPE in the absence and presence of chronic stress (Figure 1). The interaction term provides information about the genotype-dependent response to the CSE treatment. Lipidomic analysis revealed 117 and 913 lipid species that were significantly altered by either CFH genotype or CSE treatment, respectively, with 23 of the identified lipid species being affected by interaction of genotype and treatment (Figure 1A). Proteomic analysis identified 172 proteins that were significantly altered by CFH genotype alone and 7045 affected by CSE treatment, with 310 proteins affected by their interaction (Figure 1B). As shown in the Venn diagrams, the CSE treatment had a substantial effect on both lipid and protein profiles. To validate whether the lipids and proteins selected by RM two-way ANOVA can differentiate HR and LR groups, principal component analysis (PCA) was performed using the altered lipids and proteins affected by genotype or interaction. Lipids or proteins affected by the CFH genotype differentiate HR cells from LR cells under both basal (DMSO control) and CSE treatment conditions (Figure 1C, D). Lipids or proteins with significant interaction can differentiate the responses of HR from the LR groups when cells were treated with CSE (Figure 1E, F). The clear separation of groups shown by the PCA plots supports the use of RM two-way ANOVA as an appropriate method to select features for downstream analysis.

**Figure 1.**
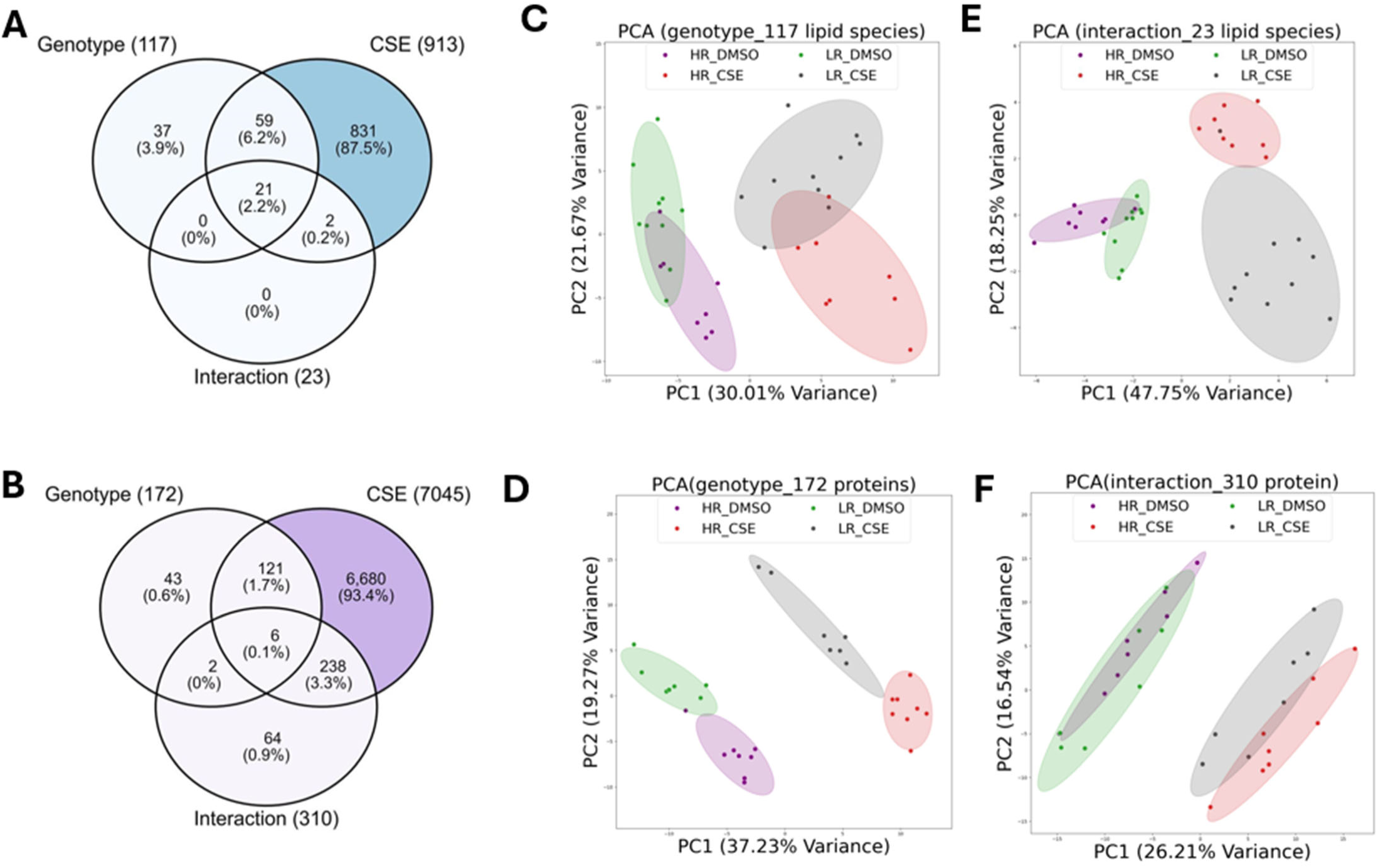
Lipid and protein profiles are affected by both CFH genotype and CSE treatment. Venn diagrams showing the number of lipid species (A) or proteins (B) altered between the CFH 402 high and low risk genotypes, by chronic CSE treatment, and the interaction of genotype and CSE treatment. Principal component analysis (PCA) was conducted using lipid species (C, E) or proteins (D, F) that were significantly altered by CFH genotype (C, D) or due to the interaction of genotype and CSE (E, F). Group separation validates the feature selection using a repeated measure (RM) two-way ANOVA.

### CSE treatment induces a significant increase in lipid biosynthesis and alters lipid subclasses

For the comprehensive lipidomics analysis, total lipids, 17 lipid subclasses (including multiple glycerolipids, phospholipids, sterols, sphingolipids, and free fatty acids) and multiple lipid species (defined by their unique structure) were identified and quantified across the different experimental groups (Figure 2A). Under basal conditions (DMSO control), total lipid content was similar in LR and HR iPSC-RPE (Figure 2B). However, treatment of the cells with CSE for two weeks significantly increased the lipid content of both LR and HR genotypes (Figure 2B), suggesting that CSE exposure significantly changes iPSC-RPE lipid metabolism. Examination of the lipid subclasses present in both genotypes found the two most abundant were phosphatidylcholine (PC) and phosphatidylethanolamine (PE), accounting for nearly 40% and 30% of total lipids, respectively (Figure 2C). These results are consistent with a prior report that PC and PE are the two most abundant phospholipids in eukaryotic membranes (23, 24).

**Figure 2.**
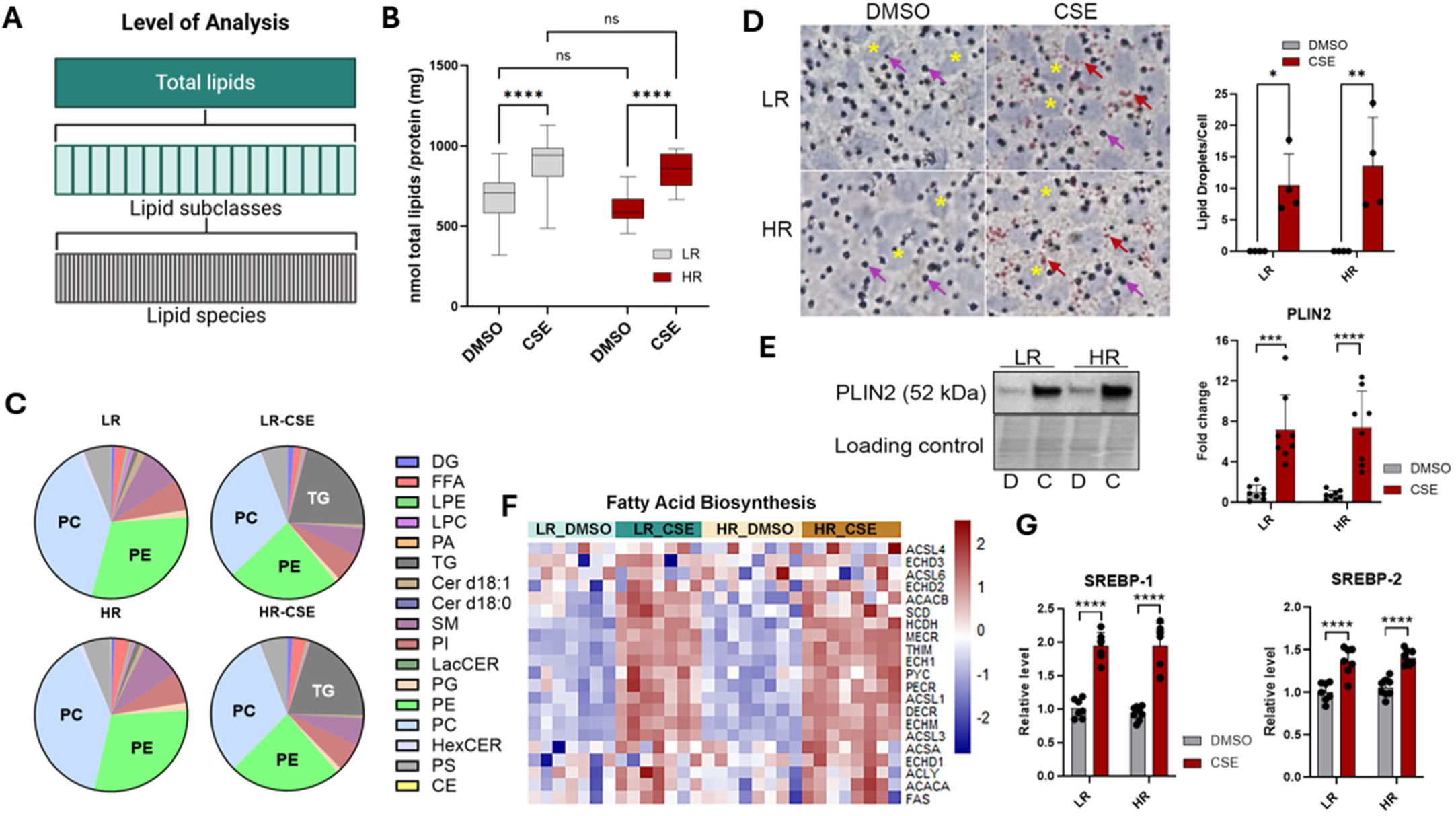
CSE treatment increases lipid biosynthesis and lipid droplet formation. (A) Diagram showing three levels of analysis in this study: total lipids, lipids from each subclass, and individual lipid species that were analyzed across all experimental groups. (B) Total lipid content quantified across groups showed CSE-treated cells exhibited significantly higher total lipid levels compared to vehicle (DMSO) controls in both CFH low-risk (LR, n=10) and high-risk (HR, n=8) cells. (C) Analysis of lipid composition at the subclass level revealed a marked increase in TG content following CSE treatment. DG (Diacylglycerol); FFA (Free Fatty Acids); LPE (Lysophosphatidylethanolamine); LPC (Lysophosphatidylcholine); PA (Phosphatidic acid); TG (Triacylglyceride); Cer d18:1(Ceramides); Cer d18:0 (Dihydroceramides); SM (Sphingomyelin); PI (Phosphatidylinositol); LacCER (Lactosylceramide; PG (Phosphatidylglycerol); PE (Phosphatidylethanolamine); PC (Phosphatidylcholine); HexCER (Hexosylceramides);); PS (Phosphatidylserine); CE (Cholesterol Esters). (D) Oil Red O staining and quantification demonstrated substantial accumulation of lipid droplets in response to CSE treatment in both LR and HR cells. n = 4/genotype. (E) Western blot and densitometric analysis of PLIN2, a lipid droplet marker, show increased PLIN2 content in CSE-treated cells. n = 8/genotype. (F) Heatmap from proteomic analysis showed significant upregulation of fatty acid biosynthesis–related proteins (gene list from WP_FATTY_ACID_BIOSYNTHESIS.v2024.1.Hs) in CSE-treated cells for both LR and HR genotypes. Z-score normalization was applied for heatmap visualization. (G) Relative SREBP-1 and SREBP-2 intensities from proteomic analysis show increased content with CSE treatment. n = 7 LR, 8 HR. Dots associated with each bar represent cell lines from individual donors. Data = mean ± SD. *p<0.05, **p<0.01, ***p<0.001, ****p<0.0001, determined from RM two-way ANOVA and Fisher’s LSD test.

Treatment with CSE, which caused a significant 30% to 40% increase in total lipid content in both genotypes, was mainly due to a substantial increase in triacylglycerides (TG) in both LR and HR cells (Figure 2B, 2C, 3A). The increased TG levels in the iPSC-RPE were confirmed by staining cells with the neutral lipid marker, oil red O (Figure 2D). Our analysis showed CSE treatment significantly increased the number of lipid droplets, which paralleled the increased expression of the lipid droplet marker Perilipin 2 (PLIN2) (Figure 2D, E). TGs are the main storage form of fatty acids, suggesting that CSE either induced *de novo* fatty acid biosynthesis or inhibited fatty acid catabolism. To gain insight into how CSE contributes to increased lipid accumulation in iPSC-RPE, the proteomics data were interrogated for content of lipid biosynthesis proteins. The data revealed significant upregulation of key proteins involved in fatty acid biosynthesis, particularly SREBP1 and SREBP2, which are key transcription factors that regulate fatty acid biosynthesis (Figure 2F, G). Further interrogation of the proteomic data showed that the majority of proteins involved in lipid biosynthetic processes were also significantly upregulated, indicating that CSE exposure triggers enhanced lipid biosynthesis (Supplement Figure 1A-B).

In addition to increased TG levels, CSE treatment significantly increased the levels of diacylglycerol (DG), phosphatidic acid (PA), phosphatidylinositol (PI), phosphatidylserine (PS), and dihydroceramide (Cer d18:0) (Figure 3A). In contrast, the levels of lysophosphatidylcholine (LPC), lactosylceramide (LacCER), hexosylceramide (HexCER), phosphatidylglycerol (PG), and cholesterol esters (CE) were significantly decreased in CSE-treated iPSC-RPE (Figure 3A).

**Figure 3.**
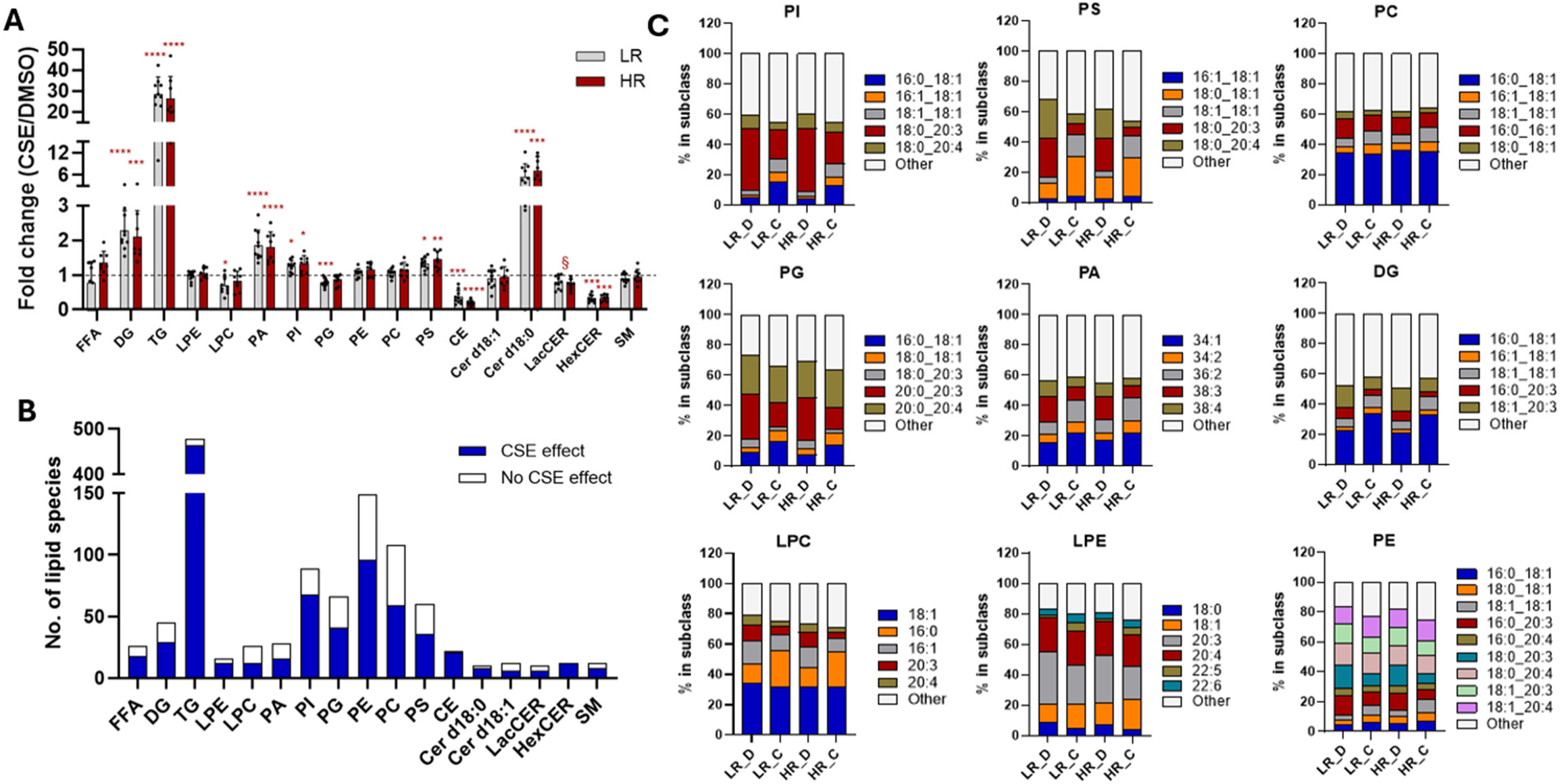
CSE treatment induces significant alterations in lipid subclasses. (A) Incubation with CSE led to changes in the levels of 10 out of 17 lipid subclasses. “§” indicates a significant CSE effect on combined LR and HR cells but not separately. Analysis of subclass levels was done using RM two-way ANOVA and Fisher’s LSD test. *p<0.05, **p<0.01, ***p<0.001, ****p<0.0001. Data = mean ± SD. (B) Number of lipid species significantly altered by CSE treatment within each lipid subclass. (C) Composition analysis showing the changes in proportion of dominant lipid species within phospholipid subclasses and DG.

Further examination of the number of lipid species affected by CSE in each subclass shows that CSE has a strong, broad effect, with some lipid species in every subclass exhibiting significant CSE-dependent changes (Figure 3B). These data suggested that the composition of lipid species within each subclass was significantly altered by CSE treatment.

To further explore how CSE affected the composition of each lipid subclass, we focused on the dominant lipid species within the phospholipids and DG subclasses (Figure 3C). The data provided shows the proportions of dominant lipid species, described by the number of carbons and double bonds for each of the two fatty acyl chains (for example DG 16:0_18:1) in DG and the phospholipids PI, PS, PC, PG, and PE. We also determined the composition of LPC and LPE, which have only one fatty acyl chain. Analysis showed that lipids containing 20:3 or 20:4 are highly abundant in PG, LPE, and in the inner-leaflet-enriched PI, PS, and PE (Figure 3C). The proportion of these species, for example PI 18:0_20:3 and PI 18:0_20:4, is significantly decreased by CSE treatment (Figure 3C). Conversely, the proportion of C16-C18 lipids with saturated or monounsaturated fatty acids (16:0, 16:1, 18:0, 18:1) in DG and most phospholipids was significantly increased by CSE (Figure 3C). Note that we were unable to identify the two specific fatty acids in PA; only the total number of carbon atoms and double bonds from the two fatty acids was determined (Figure 3C). However, we speculate that PA 38:3 and PA 38:4, which had decreased proportion, could be PA 18:0_20:3 and PA 18:0_20:4 or PA 18:1_20:3. Taken together, the data show that CSE treatment significantly decreased polyunsaturated fatty acids (PUFA) in all phospholipid subclasses and therefore altered their lipid composition.

### CSE treatment alters PUFA metabolism

Having observed a reduction in 20:3 and 20:4 after CSE treatment, we quantified the total content of different fatty acids in each phospholipid subclass and identified several fatty acids that were significantly reduced. In addition to 20:3 and 20:4, species 18:3, 18:0, and 20:0 were significantly reduced in LPC and LPE, PC, PE, PG, PS, and PI (Figure 4A, B). Although 18:0 and 20:0 are not PUFAs, they are the primary fatty acids that pair with 20:3 or 20:4 in PI, PS, PG, and PE (Figure 3C). Under our experimental conditions, the 18:0 and 20:0 likely occupy the *sn-1* position, while 20:3 and 20:4 are found in the *sn-2* position (25). This arrangement provides optimal packing order of membrane lipids(25). Lipid species significantly altered by CSE and containing any of these fatty acids (18:3, 18:0, 20:0, 20:3, 20:4) were displayed in a heatmap for each phospholipid subclass, depicting their relative abundance before and after CSE treatment (Figure 4C). After hierarchical clustering, the heat map shows a high degree of variability among lipid species before and after CSE. The checkerboard pattern reflects CSE’s complex effect on lipids, resulting in significant remodeling of the cellular lipid content and composition. Note that the most abundant lipid species in each phospholipid class, indicated in purple on the left side of each map, is decreased with CSE, and all include 20:3 and 20:4.

**Figure 4.**
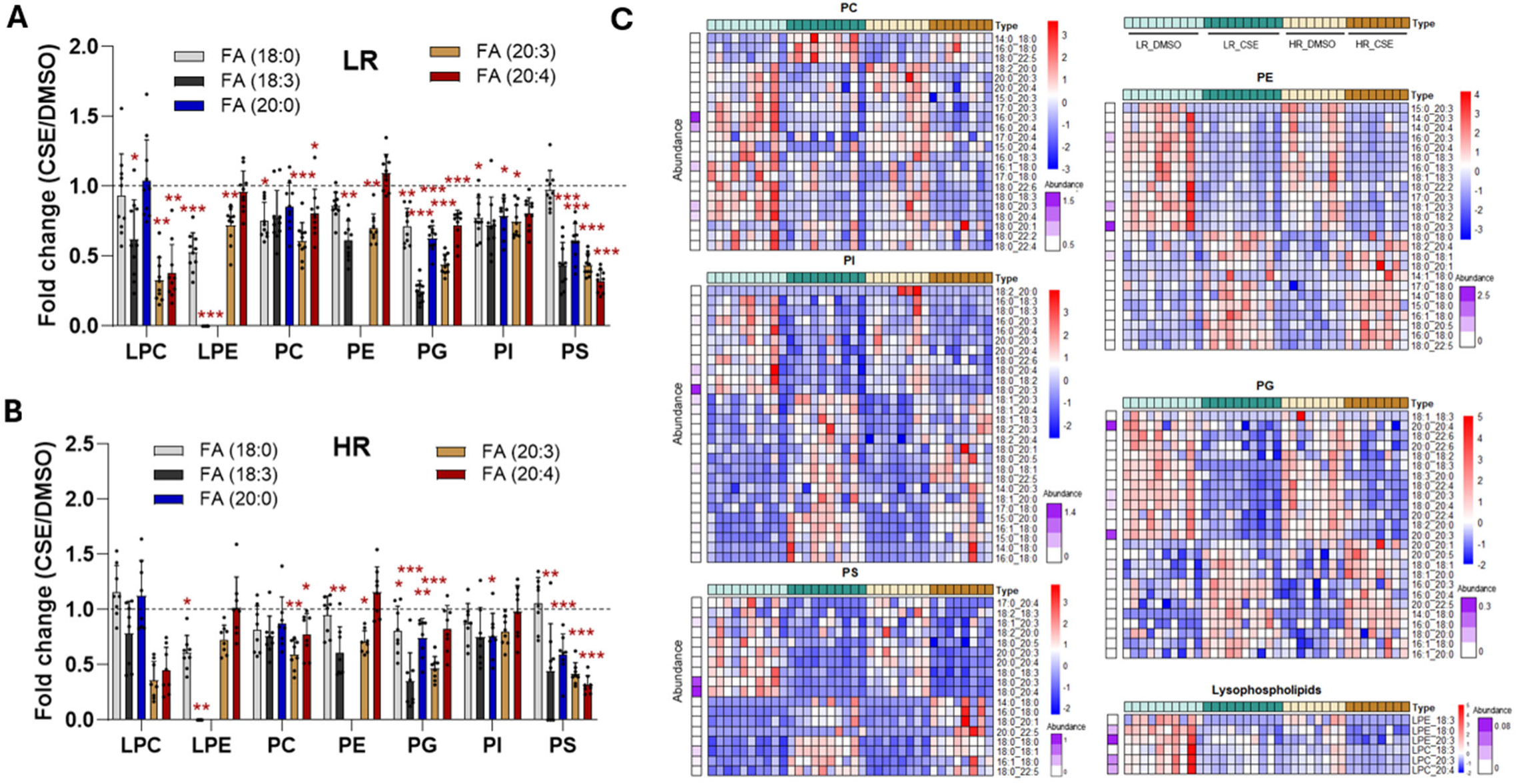
CSE treatment alters the levels of polyunsaturated fatty acids (PUFAs) in RPE phospholipids. (A, B) Bar graphs show the levels of FA 18:0, FA 18:3, FA 20:0, FA 20:3, and FA 20:4 in each lipid subclass in CSE-treated cells relative to control cells. These fatty acids were significantly reduced in most phospholipid subclasses in both CFH LR (A) and HR (B) cells following CSE treatment. Note that FA20:0 was undetectable in LPE and PE. Analysis of each fatty acid level was done using RM two-way ANOVA and Fisher’s LSD test. *p<0.05, **p<0.01, ***p<0.001. (C) Heatmaps show the normalized levels of lipid species containing selected fatty acids (FA 18:0, FA 18:3, FA 20:0, FA 20:3, and FA 20:4) in phospholipid subclasses that were significantly altered by CSE treatment. The color bar on the left (purple) reflects the abundance of each lipid species (nmol lipids/100 <g protein) in one representative sample from the LR-DMSO group.

Since the lipidomics data suggest significant remodeling of the phospholipid composition with CSE treatment, we analyzed the proteomics data to determine how CSE was affecting PUFA levels. The proteomics analyses showed that elongation of very long fatty acids 5 (ELOVL5) and fatty acid desaturases 1 (FADS1; Δ5 desaturase) and FADS2 (Δ6 desaturase), which are rate-limiting enzymes involved in PUFA biosynthesis, were significantly altered in CSE treated cells (Figure 5A-B). Dihomo-γ-linolenic acid (DGLA, 20:3) is a 20-carbon ω-6 PUFA, derived from *in vivo* elongation (by ELOVL2 and ELOVL5) and desaturation (by FADS2) of the essential fatty acid, linolenic acid (18:2n6). Subsequent desaturation of DGLA by desaturases FADS1 results in biosynthesis of arachidonic acid (AA, 20:4), another 20-carbon ω-6 PUFA (Figure 5A).

**Figure 5.**
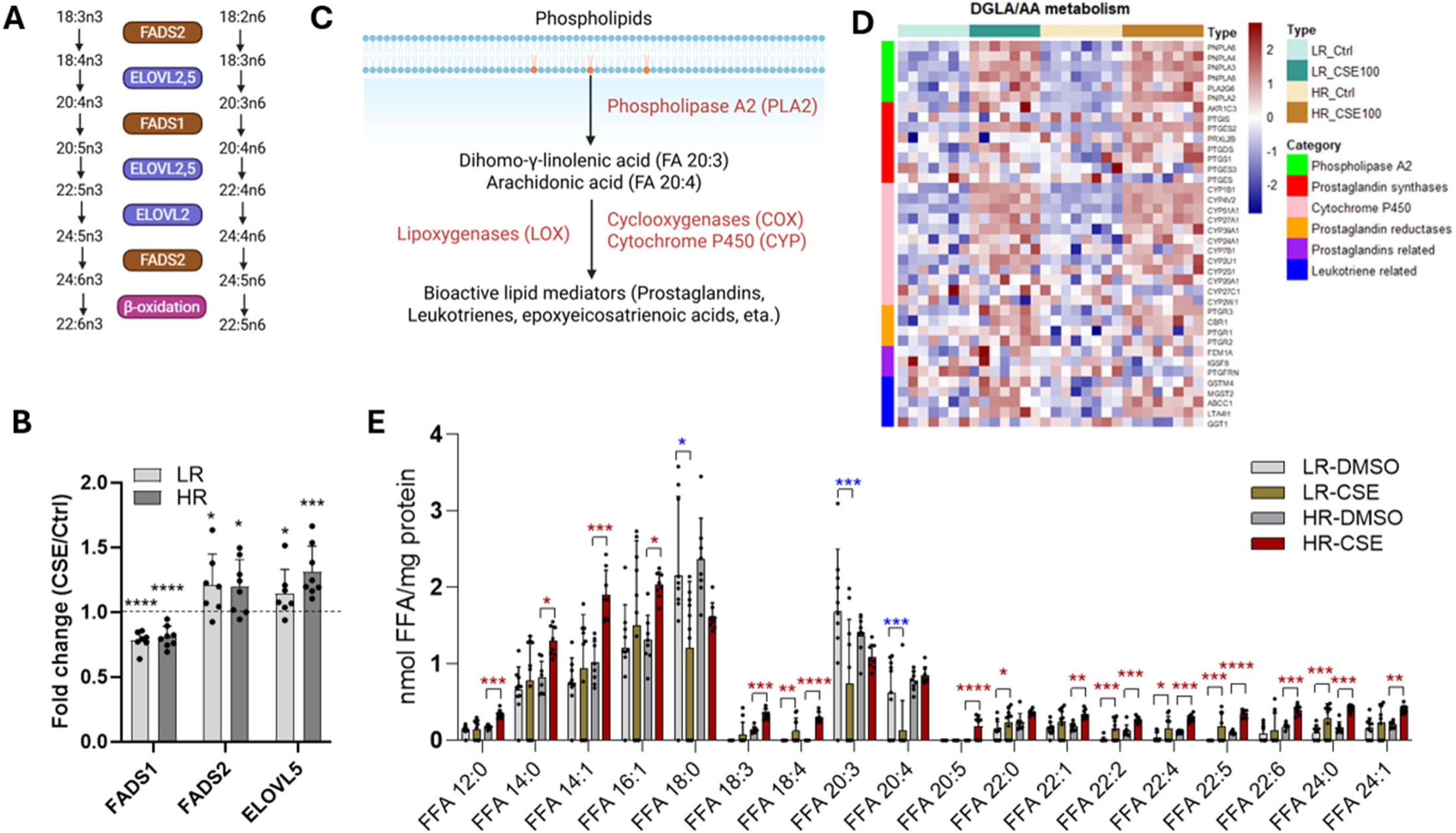
CSE treatment activates PUFA metabolism. (A) Diagram showing enzymes involved in PUFA metabolism. ELOVL, elongation of very long-chain fatty acids; FADS, fatty acid desaturase. (B) Relative level of FADS1, FADS2, and ELOVL5 in treated cells compared to DMSO controls in both LR (n=7) and HR (n=8) cells based on proteomics data. (C) Diagram showing enzymes involved in Dihomo-γ-linolenic acid (DGLA) and arachidonic acid (AA) metabolism in phospholipids. (D) Heatmap showing differential content of enzymes involved in the DGLA and AA metabolism across all groups. Color bar on the left side indicates different categories of enzymes. (E) Levels of free fatty acids significantly altered by CSE in LR (n=10) and HR (n=8) groups with DMSO or CSE. Analysis of each lipid species was done using RM two-way ANOVA and Fisher’s LSD test. *p<0.05, **p<0.01, ***p<0.001, ****p<0.0001. blue stars = decreased with CSE; red stars = increased with CSE.

Phospholipids with either DGLA or AA can be hydrolyzed by phospholipase A2 to release 20:3 and 20:4 as free fatty acids (FFA) that are further oxygenated by cyclooxygenases, lipoxygenases, and cytochrome P450 to bioactive lipid mediators, such as prostaglandins, leukotrienes, and protectins which can act as pro- or anti-inflammatory molecules (Figure 5C). Our proteomic data confirmed that the majority of enzymes involved in the hydrolysis and production of bioactive lipid mediators were significantly upregulated in CSE treated cells (Figure 5D). The release and metabolism of DGLA or AA to generate bioactive lipids could partially explain the reduced content of 20:3 and 20:4 in most phospholipids.

We also quantified the levels of FFA, which could be derived from hydrolysis not only of the fatty acyl chains in membrane lipids by phospholipase A2, but also from the hydrolysis of acyl chains from DG and TG or from *de novo* biosynthesis. Overall, there was considerable variability between the genotypes in their responses to CSE. Unique to HR cells, nine FFAs were increased; unique to LR cells, three FFAs were decreased (Figure 5E). Most importantly, FFA 20:3 and FFA 20:4 were significantly decreased in LR cells following CSE treatment, but not in HR cells. In addition, the content of some FFAs, such as 18:3, 18:4, 20:5, and 22:5, were induced by CSE as they were not detectable in DMSO control cells but were readily detected in treated cells (Figure 5E).

### CSE alters sphingolipids and cholesterol content

Two other subclasses linked to AMD pathology, cholesterol and sphingolipids, were also significantly altered by CSE treatment. Ceramide metabolism involves multiple, reversible biosynthetic pathways (Supplement Figure 2A). Ceramide (Cer d18:1) can be synthesized *de novo* using L-Serine and palmitoyl-CoA, with the dihydroceramide (Cer d18:0) as the precursor. It can also be generated from the degradation of glucosylceramides, such as Hexosylceramides (HexCER) and Lactosylceramide (LacCER) through the salvage pathway or by the hydrolysis of sphingomyelins (SM). We quantified ceramide, dihydroceramide, sphingomyelin (SM), lactosylceramide (HexCER), and hexosylceramide (HexCER) across all four experimental groups. Dihydroceramides (Cer d18:0) were increased while HexCER and LacCER weredecreased significantly in cells after CSE (Figure 3A), indicating that the treated cells have been able to increase the ceramide pool. However, Cer d18:1 (ceramide) level is not significantly altered by CSE, suggesting that it is consumed for other purposes. Proteomic data showed significant alterations in the content of enzymes involved in ceramide metabolism, especially ASAH1, an enzyme responsible for converting ceramide to sphingosine, that had a two-fold increase in treated cells. This observation, together with the increase of enzymes for sphingosine-1-phosphate synthesis and breakdown (SPHK2 and SGPP1), suggested an active ceramide to sphingosine-1-phosphate pathway (Supplement Figure 2A).

Cholesterol was also highly altered by CSE treatment. Lipidomic analysis showed that nearly all cholesterol ester (CE) species were significantly decreased, with some species becoming undetectable after 2 weeks of CSE treatment (Supplement Figure 2B). Consistent with the reduced levels of CE, we observed that the expression of enzymes involved in CE hydrolysis (NCEH1, LIPE, LIPA, and CES1) and free cholesterol transporters, ABCA1 and ABCG1, were significantly upregulated by CSE treatment (Supplement Figure 2C). In contrast, SOAT1, the enzyme which esterifies excess intracellular cholesterol for storage, was not changed by CSE but had a significantly lower content in HR. These data strongly suggest enhanced hydrolysis of CE and cholesterol efflux from the cell under stressed conditions.

### CFH HR RPE exhibit higher levels of free fatty acids before and after CSE treatment

In addition to the dramatic effects of CSE on RPE lipid and protein profiles, the CFH genotype had a significant influence on the lipid and protein profiles before and after CSE treatment. Our analysis showed that 119 lipid species were affected by either genotype or by the interaction between CFH genotype and CSE treatment (Figure 1A, Supplement Figure 3). At the subclass level, HR cells had higher FFA and CE but lower LPE, LPC, and DG, either at basal levels or under CSE treatment (Figure 6A). Under basal conditions, 14 species were higher, including 5 FFAs and 5 CEs, and 17 were lower, including 4 LPE and 6 LPC, in HR compared with LR iPSC-RPE (Figure 6B). Two FFAs (18:3 and 22:5) and PI 18:2_20:0 were uniquely present in only HR cells. Under CSE-induced stress, HR cells exhibited increases in 39 lipid species, while 26 were decreased. Of note, FFA 20:0, 20:5, and PC 20:0_20:1 were only detected in HR cells.

**Figure 6.**
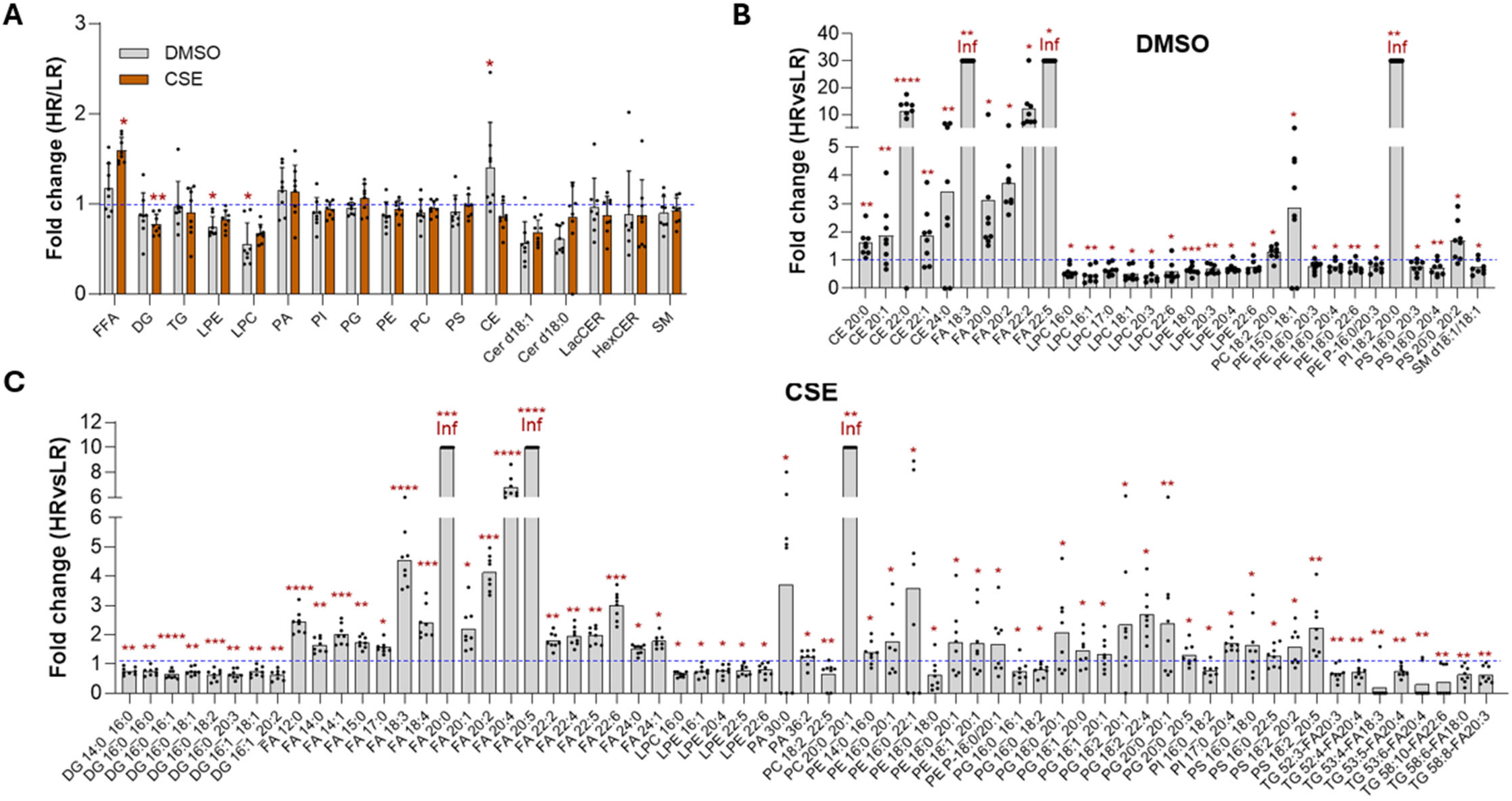
CFH genotype affects free fatty acid levels under basal and CSE-treated conditions. (A) Bar graphs depict relative levels of each lipid subclass in HR cells compared to LR cells under basal (DMSO control) and CSE-treated conditions. (B-C) Bar plots show fold change for lipid species exhibiting significant differences between HR and LR cells under basal (B) and CSE-treated (C) conditions. Lipids with no detectable signal in the LR group (resulting in fold change = ∞) are marked as “Inf.” For visualization, bars were capped at fold change = 30 (B) and 10 (C), respectively. Significance was generated using RM two-way ANOVA and Fisher’s LSD test. *p<0.05, **p<0.01, ***p<0.001, ****p<0.0001.

Additionally, there were increases in FFA 18:3, 20:4, 22:5, and 22:6, and corresponding lower levels of TGs with 18:3, 20:4, 22:6, and LPEs with 20:4, 22:5, and 22:6 (Figure 6C). These results show prominent targeting of PUFA hydrolysis with CSE and suggest that lipid hydrolysis may be dysregulated in HR cells. It is worth noting that 29 lipid species did not show significant differences in either condition in the post hoc analysis, but they did show a significant genotype effect in the RM two-way ANOVA (Supplement Figure 3).

### CFH HR RPE exhibit higher levels of lipolytic proteins and altered inflammation proteins

Analysis of proteomics data using PCA for the 200 proteins that show differential expression after CSE treatment demonstrates excellent reliability in separating the HR and LR groups (Figure 7A). Among the 200 differentially expressed proteins, 28 proteins are related to lipid handling (Figure 7B), including abhydrolase domain containing 5 (ABHD5) and patatin-like domain 2, triacylglycerol lipase (PNPLA2, or ATGL), two proteins that are directly involved in TG hydrolysis. ABHD5 directly interacts with and activates ATGL, which hydrolyzes TG (26). In support of the proteomics results, Western blot analysis confirmed a higher level of ABHD5 in HR cells compared with LR cells after CSE treatment (Figure 7C). Although the total TG content was not significantly different in HR cells after CSE treatment (Figure 6A), the levels of PUFAs in their storage form (TG) and FFA form were altered between genotypes (Figure 6C). In HR cells, levels of specific TG species were significantly reduced whereas specific FFA species were significantly increased compared with the response in LR cells. As previously mentioned, PUFA are precursors for bioactive lipid mediators that regulate inflammation and participate in the immune response (Figure 5A, C). Consistent with the genotype specific differences in precursors of bioactive lipids, our proteomics data also show a genotype-dependent response to CSE for proteins involved in regulation of inflammation (Figure 7D).

**Figure 7.**
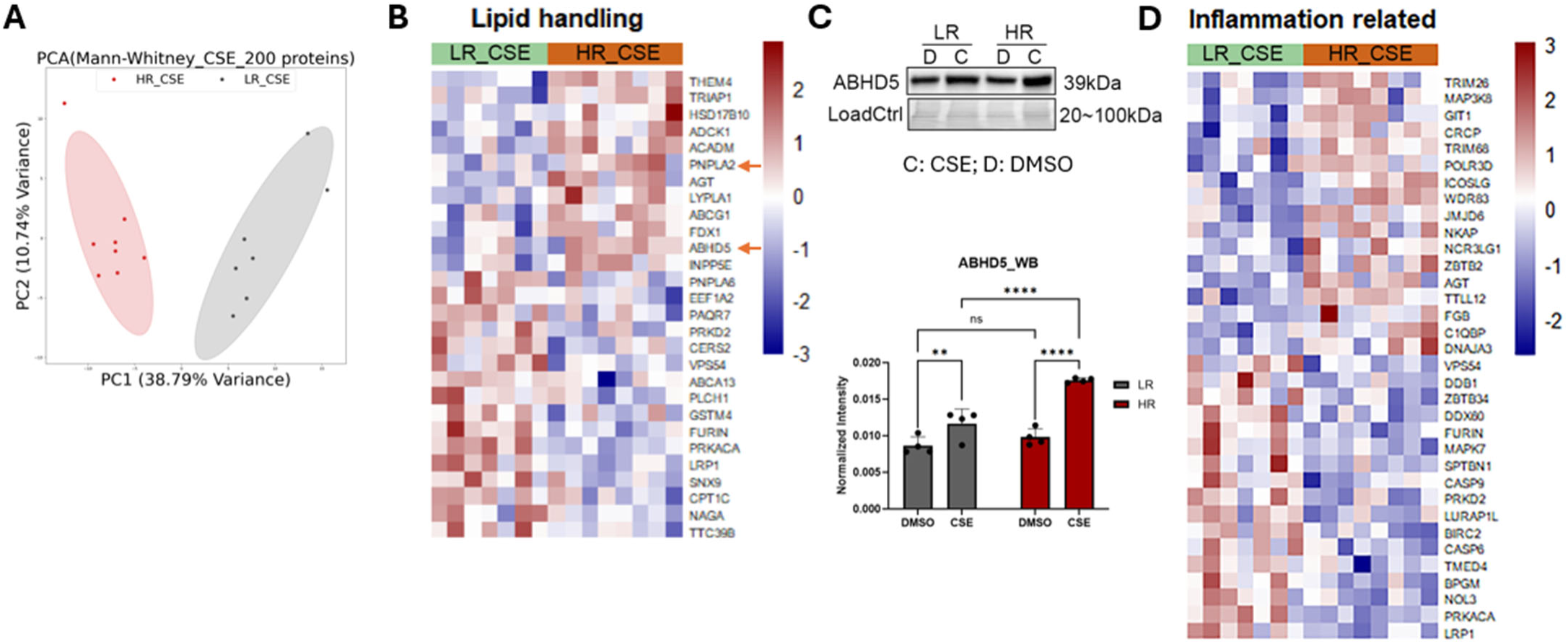
CFH HR RPE exhibits elevated levels of lipolytic proteins after CSE treatment. (A) Principal component analysis (PCA) was conducted using altered proteins identified by Mann Whitney test comparing HR cells (n=8 donor lines) with LR cells (n=7 donor lines) following CSE treatment. (B) Proteins involved in lipid handling were significantly altered in HR cells compared to LR cells after CSE. (C) Western blot and results of densitometric analysis showed increased ABHD5 in HR cells compared to LR cells after CSE exposure. Data = mean ± SD. **p< 0.01, ****p<0.0001, ns = no significant difference. (D) Content of proteins involved in inflammatory processes were significantly altered in HR cells compared to LR cells by CSE.

### CFH HR RPE has lower fatty acid oxidation

The FFA pool in a cell is highly dynamic and is regulated by different processes. For example, FFAs can be generated by *de novo* fatty acid biosynthesis or by their release from membrane lipids via hydrolysis. Alternatively, FFA content can be reduced when they are incorporated into different lipid subclasses or serve as substrates for fatty acid oxidation (FAO), a process that generates energy in the mitochondria. We have previously shown that CFH HR iPSC-RPE had more significant mitochondrial dysfunction compared to LR iPSC-RPE following CSE treatment (17). Considering that FFAs are generally higher in HR cells both before and after CSE, we hypothesized that FAO may be reduced in HR cells. To test this idea, we performed a Fatty Acid Stress Test on LR and HR iPSC-RPE treated with either DMSO (vehicle control) or CSE for one week. FAO is determined by measuring the oxygen consumption rate (OCR) in iPSC-RPE cultures in the absence and presence of etomoxir, an irreversible inhibitor of carnitine palmitoyltransferase 1a (CPT1a), which transports fatty acids into mitochondria (Figure 8A).

**Figure 8.**
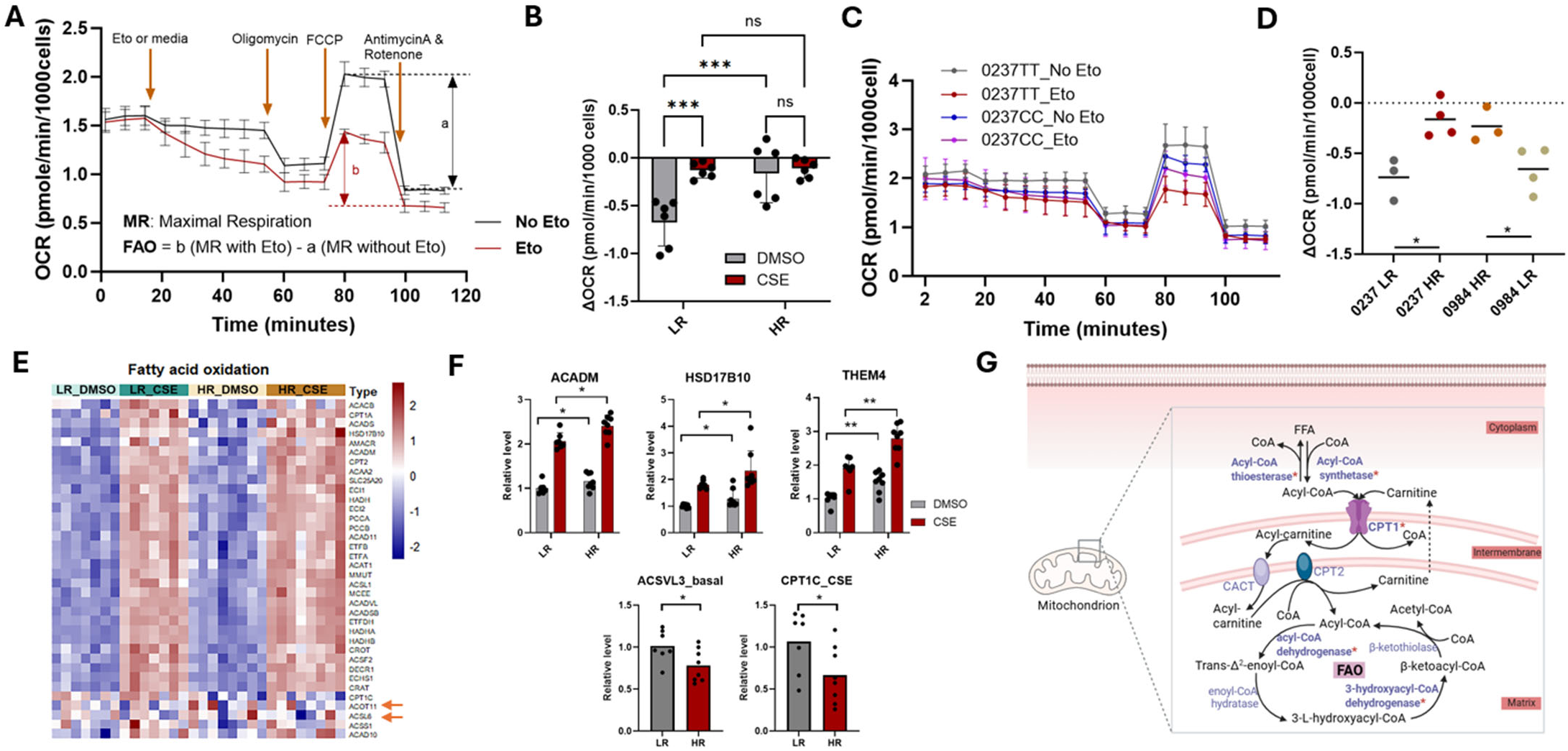
CFH HR RPE exhibit lower fatty acid oxidation. (A) Trace of oxygen consumption rate (OCR) during a Fatty Acid Oxidation Stress Test shows the method for calculating the level of cellular fatty acid oxidation (FAO). Eto (etomoxir) is an FAO inhibitor. (B) Quantification of FAO in CFH LR and HR cells with or without CSE treatment. n = 6/genotype, (C) Traces of OCR during an FAO assay for a parent line (0237 LR) and the corresponding isogenic line (0237 HR), with and without the FAO inhibitor, etomoxir (Eto). (D) FAO levels quantified in two parent lines and their corresponding gene-edited isogenic lines resulting in an amino acid change at position 402 in the CFH protein. n = technical repeats. (E) Heatmap showing relative levels of proteins involved in fatty acid oxidation (Mito Carta 3.0) across groups. (F) Bar graphs showing genotype differences in proteins related to FAO. ACADM (medium-chain specific acyl-CoA dehydrogenase), HSD17B10 (3-hydroxyacyl-CoA dehydrogenase type-2), THEM4 (Acyl-CoA thioesterase THEM4), ACSVL3 (very long chain fatty acyl-CoA synthetase 3), CPT1C (carnitine O-palmitoyltransferase 1C). Analysis was done using RM two-way ANOVA and Fisher’s LSD test. (G) Figure illustrates the FAO pathway and function of the associated proteins. Proteins with genotype-dependent differences in content are indicated with “*” in the FAO pathway. Data = mean ± SD. *p<0.05, **p< 0.01, ***p< 0.001, ns = no significant difference.

Under basal conditions, HR cells exhibited significantly lower FAO compared to LR iPSC-RPE (Figure 8B). Treatment with CSE nearly eliminated FAO in both LR and HR cells (Figure 8B). Due to the overwhelming effect of CSE on FAO, no genotype-dependent effect could be differentiated. To confirm the genotype-dependent difference in FAO under basal conditions, FAO was measured using pairs of parent and their isogenic iPSC-RPE cell lines in which the CFH Y402H risk SNP was edited from LR to HR or vice versa. We generated iPSC-RPE from isogenic edited iPSC lines from two different donors. Parent line 0237 is LR, and the corresponding isogenic line is edited to HR. Parent line 0984 is HR, and the isogenic line is edited to LR. To determine how the change in a single amino acid at position 402 in the CFH protein affects function, we compared FAO between the parent line and their corresponding isogenic line, which have the same genetic background except for the edited allele. Figure 8C shows OCR during an FAO assay for the 0237 LR parent line and the corresponding isogenic edited 0237 HR line with and without the FAO inhibitor, etomoxir (Eto). In these two pairs of parents and isogenic edited iPSC-RPE lines, the HR parent or HR edited iPSC-RPE showed reduced FAO compared to the corresponding LR lines (Figure 8D). These results agree with our initial findings using iPSC-RPE from multiple individual donors showing that cells harboring the CFH HR genotype have lower FAO. Reduced utilization of FFA as a substrate for FAO could partially account for the increase in FFA levels observed in HR cells.

Even though CSE treatment strongly suppressed FAO, proteomics data show that CSE elicited a substantial increase in the majority of FAO proteins (protein list from Mito carta 3.0), except for ACOT11 (acyl-CoA thioesterase 11) and ACSL6 (acyl-CoA synthetase long chain family member 6), which were either not changed (ACOT11) or decreased (ACSL6) by CSE treatment (Figure 8E). Taken together, the proteomic results suggest that, though the cells have initiated a compensatory mechanism in an attempt to rescue FAO by increasing the machinery proteins, the CSE treated cells still had lower FAO rates. Proteomics data also revealed significant genotype differences in proteins related to FAO (Figure 8F). ACADM (medium-chain specific acyl-CoA dehydrogenase), HSD17B10 (3-hydroxyacyl-CoA dehydrogenase type-2), and THEM4 (Acyl-CoA thioesterase THEM4) exhibited elevated content in HR cells compared to LR cells under both basal and CSE conditions. Conversely, ACSVL3 (very long chain fatty acyl-CoA synthetase 3) and CPT1C (carnitine O-palmitoyltransferase 1C) showed reduced content in HR cells under basal and CSE conditions, respectively. The bidirectional pathway regulated by THEM4 and ACSVL3 directly contributes to cellular levels of FFA and Acyl-CoA (Figure 8G). Decreased ACSVL3 and increased THEM4 could increase FFA levels. These data suggest that dysregulation of multiple processes contributes to the altered FFA content and the reduction of FAO in HR cells.

## Discussion

This study investigated how the CFH Y402H high risk SNP for AMD alters lipid homeostasis in iPSC-RPE. Using a combination of discovery-based and targeted assays, we studied iPSC-RPE generated from CFH high and low risk donors in the presence and absence of CSE stress to understand how these prominent genetic and environmental AMD risk factors contribute to an individual’s risk for AMD. Significant genotype-dependent differences in the lipid and protein profiles under basal and CSE-treatment were observed. Under basal conditions, HR iPSC-RPE exhibited higher levels of FFA and CE; CSE further amplified the differences in FFA content. HR cells also had significantly lower FAO compared with their corresponding LR line when measured in multiple donors and in pairs of parent/ isogenic edited lines. Consistent with these results, proteomic data revealed genotype specific responses to CSE-induced stress, including dysregulated lipid handling and lipolysis, and differential content of proteins involved in inflammation and fatty acid oxidation. Furthermore, we found that exposure to CSE for two weeks had a profound effect on both lipid and protein profiles in HR and LR iPSC-RPE. Data from lipidomic analysis following CSE exposure showed significant remodeling of cellular lipid content and composition, including lipid accumulation, changes in composition of lipid species for all lipid subclasses, and a significant loss of CE and PUFAs in phospholipids. Supporting these findings, proteomic analysis identified upregulation of proteins involved in lipid synthesis and hydrolysis, production of bioactive lipid mediators, and metabolism of ceramide and cholesterol. Taken together, our results show that layering AMD genetic and environmental risk factors allowed us to tease out CFH genotype-dependent differences in RPE lipid homeostasis. These novel results help to elucidate the potential causes for disease susceptibility and the underlying mechanisms driving pathology in individuals carrying the CFH risk alleles.

The goal of the combined “omics” approach was to obtain an unbiased overview of how CFH genotype and CSE treatment affect the lipid and protein profile in the RPE. The detailed lipidomic analyses revealed important new insights into potential functional changes in RPE lipid metabolism due to CFH genotype and CSE, which altered the composition of lipid subclasses and their fatty acyl chains. These fine details are important when considering the metabolic fate of outer segment- and diet-derived lipids in the RPE in individuals with the HR CFH genotype who are also smokers. RPE function requires the dynamic formation of apical processes for retinal outer segment phagocytosis and the formation of autophagosomes for the clearance of retinal outer segment membranes, which are influenced by the phospholipid composition of the cell membrane. As the major structure component of membranes, the amphipathic nature of phospholipids and their acyl chains allow cholesterol and other lipids to be incorporated into the lipid bilayer, which is essential for cellular function. The fatty acid composition of the bilayer also regulates membrane stability and fluidity. It also creates cellular compartments for biosynthesis of glycerolipids (TG, DG), that contribute to membrane structure, store energy (TG) and are involved in cell signaling (DG and other lipids). The function of transmembrane proteins, such as channels, receptors, and pumps, is highly dependent on the composition of the surrounding lipid bilayer. Thus, many of the changes in lipid composition observed in this study could have significant effects on membrane fluidity and consequently impact the important cellular functions of the RPE that are necessary to maintain vision. For example, the kinks and bends in PUFAs caused by their double bonds disrupt tight packing of saturated membrane lipids, increasing membrane fluidity and allowing proteins embedded in the membrane to move more freely.

Cholesterol has a buffering effect on fluidity by interacting with the fatty acid chains, thereby maintaining membrane stability under diverse conditions. This suggests that the CSE-induced reduction in PUFAs and near elimination of CE would greatly reduce membrane fluidity, potentially affecting the function of proteins embedded in the cell membranes (Fig. 4, Supplement Fig 2). Lipid bilayer composition is also critically important for processes involving membrane fission and fusion, as well as vesicle formation associated with exocytosis, endocytosis, and autophagosome formation. Autophagosome formation is critical for retinal health, including the transport and degradation of phagocytosed outer segments and the recycling of retinoid cycle products. The formation of an autophagosome requires phospholipids to form the double-membrane vesicles that enclose cellular material destined for degradation after fusion with lysosomes during the autophagy process (27). Our findings that the CFH genotype and CSE altered RPE lipid biosynthesis with resultant increased TG accumulation suggest that RPE autophagosome lipid membrane composition could be altered, which could impact autophagy and the RPE-specific process of outer segment degradation.

Lipids perform a second critical role as bioactive second messengers that regulate various cellular processes, such as inflammation and the immune response. For example, hydrolysis of both omega-3 and omega-6 fatty acids by phospholipase A2 can release the FFA 20:3 or 20:4, that can be further processed to bioactive lipid mediators, such as protectins, which are derived from omega-3 FA and can actively stop inflammation and promote tissue healing (28). CSE exposure had a significant effect on phospholipid PUFA levels, as evidenced by their overall reduction in content and increased content of lipolytic enzymes involved in generating bioactive lipid mediators, such as prostaglandins and leukotrienes (Fig. 5). Of note, genotype-dependent differences were observed for PUFAs in their storage form (TG) and free form (FFA), and in proteins involved in lipolysis and inflammation (Fig. 5,6,7). Other bioactive lipids include ceramides, whose specific function is dependent on their fatty acyl chain composition, and DG. As an activator of protein kinase C, DG is central to many cellular functions, including proliferation, cytokine production, and metabolism. The lower content of DG observed in untreated HR iPSC-RPE and after CSE treatment (Fig. 6) provides a potential mechanism for dysregulation of multiple cellular function regulated by protein kinase C in the RPE of AMD HR patients.

One additional critical function for RPE lipids is their use as a substrate for production of ATP via FAO through mitochondrial β-oxidation. While glucose is readily available for production of energy via glycolysis by the RPE, seminal work by the Hurley and Philp research groups have shown that glucose from the choroid is shunted through the RPE into the interphotoreceptor space and taken up by the photoreceptors, where it is used for aerobic glycolysis to produce ATP and lactate as a byproduct (29, 30). The photoreceptor-generated lactate is transported to and taken up by RPE where it is used as a substrate for oxidative phosphorylation (OxPhos). In addition to lactate, RPE can utilize a variety of substrates, including glycogen, glutamine and other amino acids, to fuel OxPhos (31). However, the most abundant fuel is lipids, which in vivo, mainly come from the daily phagocytosis and degradation of outer segments. Lipids processed to FFA are oxidized by mitochondrial FAO to generate NADH, FADH2, and acetyl CoA, which fuel OxPhos. Ketone bodies are also produced by FAO, then exported from the apical side of RPE, taken up by photoreceptors, and used as an energy source (30). Thus, there is a metabolic co-dependence between the RPE and photoreceptors that requires maintaining the flow of shared substrates for ATP production. This “metabolic ecosystem” is disrupted when RPE mitochondrial function is reduced, forcing the RPE to rely on glucose and glycolysis to generate energy, thereby eliminating production of ketone bodies for the photoreceptors (32).

The vicious cycle initiated by RPE mitochondrial dysfunction creates a bioenergetic crisis that has been proposed as one of the pathological defects driving AMD. The mechanism is well supported by reports of damaged mtDNA in RPE harvested from AMD donors, and measurements of reduced RPE mitochondrial function along with increased RPE glycolysis in cultured cells from AMD patients (16, 17, 33, 34).

Considering the high proportion of AMD patients harboring the HR CFH Y402H allele, we previously investigated the possibility that the presence of this CFH SNP was driving these results. By separating donor samples based on their genotype, we found the AMD donors carrying the HR CFH SNP had significantly more mtDNA damage than donors with LR alleles (33). In another study, analysis of iPSC-RPE derived from donors with the LR or HR CFH SNPs showed mitochondrial function was significantly reduced in HR cells, irrespective of the AMD disease stage of the donor (16). More recently, we showed that treatment of iPSC-RPE with CSE caused a dose-dependent decrease in mitochondrial function that was significantly more pronounced in the HR cells (17). Results from the current study extended and refined defects in mitochondrial function to include reduced FAO in RPE from donors with the HR CFH SNP (Fig. 8).

Additional supportive evidence was provided by testing parent iPSC-RPE compared with their isogenic lines in which the nucleotide responsible for the CFH risk SNP was edited. Since the parent and isogenic lines have identical genetic backgrounds, any difference in response can be attributed to the single amino acid change at position 402 in the CFH protein. In this comparison, cell lines with the HR genotype had significantly lower FAO, which is consistent with the result in iPSC-RPE from multiple donors. Thus, we are confident that the CFH SNP has a significant negative effect on mitochondrial FAO. This reduction in FAO could be responsible, in part, for the increase in FFA in HR iPSC-RPE under basal and CSE-treated conditions (Fig. 6). Based on the discussion above, the potential negative effect of reduced FAO on RPE could have far-ranging effects on the health and survival of the photoreceptors. Indeed, defects in FAO due to inherited FAO disorders often result in RPE retinopathy followed by photoreceptor death (35–37).

Lipid metabolism is a tightly regulated process that, when disrupted, can trigger lipid accumulation both outside and within the RPE. One of the hallmarks of AMD is the formation of lipid-rich drusen deposited beneath the RPE and throughout Bruch’s membrane, the extracellular matrix between the choroid and RPE. Drusen is made up of cholesterol, PC, and TG, which are derived from the circulation and from lipids produced within the RPE. These lipid deposits form a lipid wall, which can impede the flow of nutrients and oxygen into and waste products out of the RPE (38). This disruption of nutrients, oxygen, and waste exchange can lead to photoreceptor cell death and induce neovascularization, characteristics of the advanced stages of AMD known as geographic atrophy and neovascular AMD, respectively.

The current study showed CFH HR cells had elevated levels of cholesterol esters (CE) compared to LR cells under basal conditions (Fig. 6). Parallel proteomic analysis showed the content of the cholesterol transporter ABCG1 was higher, while there was reduced content of SOAT1, the enzyme that generates CE (Supp Fig 2). These results suggest HR cells may export more free cholesterol, which could contribute to increased drusen formation. Consistent with this idea, analysis of an AMD patient population showed that individuals with the CFH HR SNP had elevated drusen content compared with LR patients, and that the extent of drusen in HR patients correlated in an allelic dose-dependent manner (39). An additional link between drusen formation and the CFH SNP is provided by results using iPSC-RPE derived from patients genotyped for the CFH SNP. They observed an accumulation of lipid droplets and “drusen like” deposits under the RPE in cells with the CFH HR genotype (19). Taken together, dysregulation of lipid handling and altered lipid content associated with the CFH HR SNP could provide the mechanistic basis for the increased drusen formation associated with AMD.

This study employed the most advanced instruments and analytical platforms to perform in-depth quantification of lipids and proteins, resulting in identification of multiple cellular processes affected by the CFH Y402H SNP and/or CSE exposure. However, some technical limitations restricted our capacity to fully probe the molecular details of these changes. For example, one of our main interests is in understanding why mitochondrial function, including FAO, is reduced in RPE with the CFH HR SNP. The bulk analysis of lipids that was performed does not provide information on how the CFH genotype and CSE interaction affect the lipid composition of the plasma membrane and of individual organelles. Additionally, low-abundance lipids, such as the mitochondria-specific lipid cardiolipin, were outside the range of detection. Cardiolipin is localized to the inner mitochondrial membrane where it provides structural stability required for optimal function of energy-generating proteins in the electron transport chain. Another limitation is that our analysis cannot identify the position of double bonds and the fatty acyl chains in the *sn1* or *sn2* positions. This information could differentiate the omega-3 and omega-6 fatty acids, which produce either pro- or anti-inflammatory bioactive lipids. The n-3/n-6 ratio can be used to predict the cell’s inflammatory potential. Free cholesterol was not measured in the lipidomic analysis, which limited our evaluation of cholesterol metabolism changes based on data of CE from lipidomics and cholesterol transporters from proteomic analysis. A limitation of our proteomics data is that protein content alone does not accurately predict protein and enzyme function. Additional information, such as post-translational modifications or the association of cofactors, is required to gain a more complete understanding of how activity of specific pathways is affected by the CFH SNP or CSE.

Our experimental design examined the interconnection of two biologically distinct pathways linked to AMD: lipid metabolism and the complement system. While the importance of each pathway alone has been well-recognized, recent studies in humans, mice, and cell culture models have strengthened the link between abnormalities in lipid metabolism and complement system dysregulation. For example, a prospective study of more than 450,000 genetically defined subjects reported that lipid-lowering drugs protected against developing AMD in individuals carrying the HR CFH SNP. Additionally, the lipid lowering drugs reduced their rate of AMD progression by 50% (40). Transgenic mice expressing the human HR CFH protein had increased deposits of subretinal lipids and lipoproteins compared with mice expressing the LR CFH protein (41). In iPSC-RPE developed from HR CFH patients, lipid droplet accumulation and deposition of “drusen-like” deposits were observed (18). These data indicate an intricate interaction between these two crucial pathways that may act synergistically to contribute to AMD disease onset and progression.

In conclusion, this investigation used discovery-based and targeted approaches to reveal significant differences in the lipid and protein profiles in iPSC-RPE harboring either the LR or HR CFH genotype. In addition, functional measurements showed FAO was significantly reduced in iPSC-RPE with the HR genotype. These results broaden our understanding of the intracellular role for CFH in an expanding list of processes, such as regulating multiple degradative pathways, that are essential for RPE function and survival.

## Methods

### Sex as a biological variable

Our study examined cells from male and female donors, and similar findings are reported for both sexes.

### iPSC-RPE cell culture and CSE treatment

The derivation of iPSC from epithelial cells harvested from the conjunctiva of human donor eyes and their differentiation into RPE cells was performed as previously described (42). Cell culture conditions were followed according to established protocols (16, 17, 42). iPSC-RPE monolayers from passages 3 or 4 were cultured for two weeks to allow sufficient time for cells to become confluent before starting CSE treatment. CSE (NC1560725, Murty Pharmaceuticals, Lexington, KY) was purchased as a stock solution (40 mg/ml) in DMSO and stored at 4 ° C. CSE (final concentration 100 μg/mL) was administered during media changes three times per week for two weeks. DMSO served as vehicle control. Samples were collected 48 h after the last treatment for downstream analysis.

### Isogenic iPSC-RPE generation

Generation of the isogenic lines used in this study was accomplished by editing the CFH gene in iPS cells derived from human donor eyes using CRISPR-Cas9 technology. Guide RNAs were designed including the one nucleobase switch in the CFH gene (nucleotide position 1277 in exon 9) exactly before a PAM (Protospacer Adjacent Motif, TGG in this CFH gene) sequence and electroporated into iPS cells together with CAS9 and introduction of HDR template using AAV6 vectors. Single cells were cultured at low density and colonies with CFH edits were identified by DNA sequencing. CFH edited lines were cultured and expanded, then differentiated into RPE. Guide and template sequences used are:

CFH Y402 (donor 1, TT to CC) gRNA - 5’-AAUGGAUAUAAUCAAAAUUA-3’ DNA Version - 5’-AATGGATATAATCAAAATTA-3’.

CFH H402 (donor2, CC to TT) gRNA - 5’-AAUGGAUAUAAUCAAAAUCA-3’ DNA Version - 5’-AATGGATATAATCAAAATCA-3’.

### Shotgun lipidomics

Shotgun lipidomics was performed at the UCLA Lipidomics core facility (Los Angeles, CA). iPSC-RPE cells were cultured in 12-well plates and treated with either DMSO or CSE (100 <g/mL) three times per week during media changes. After two weeks of treatment, cells were scraped, pelleted in PBS by centrifugation, and snap-frozen as cell pellets.

Lipid extraction was performed using a modified Bligh and Dyer method (43). A standard mixture containing 74 lipid standards spanning 17 subclasses (Cat # 330820, 861809, 330726, 330729, 330727, and 791642, Avanti Polar Lipids, Alabaster, AL) was added to each sample. After two successive extractions, the pooled organic layers were dried using a Thermo SpeedVac SPD300DDA (ramp setting 4, 35 °C, 45 minutes; total run time: 90 minutes). Dried lipid extracts were resuspended in a 1:1 mixture of methanol and dichloromethane containing 10 mM ammonium acetate and transferred to Robovials (Cat. # 10800107; ThermoFisher Scientific, San Jose, CA) for analysis.

Samples were analyzed using the Sciex Lipidyzer platform (Sciex 5500 with DMS and Shimadzu LC-30) for targeted quantification of 1,400 lipid species (known abundant in mammalian tissue and cells) across 17 subclasses. The Differential Mobility Spectrometry (DMS) device was tuned using EquiSPLASH LIPIDOMIX standard (Cat # 330731; Avanti Polar Lipids, Alabaster, AL). Instrument parameters, tuning conditions, and the multiple reaction monitoring (MRM) list have been previously published (44). Data analysis was conducted using Shotgun Lipidomics Assistant (SLA v1.3) (44). Quantitative lipid values were normalized to protein concentrations derived from parallel duplicate wells.

Principal component analysis (PCA) and heatmaps were generated to assess the global structure of the dataset and examine variable relationships. Outliers identified via PCA were excluded from the final analysis. Total lipids, lipid subclass totals, and individual lipid species were analyzed between groups using repeated measures two-way ANOVA and Fisher’s LSD test.

### Protein preparation and label-free mass spectrometry

After two weeks of CSE treatment, cells were scraped, pelleted in PBS by centrifugation, and snap-frozen as cell pellets. For protein extraction and digestion, an SC-aided precipitation/on-pellet digestion (SEPOD) protocol was employed as previously described (45). Protein was first reduced by 10 mM dithiothreitol (DTT) at 56 °C for 30 min and alkylated by 25 mM iodoacetamide (IAM) at 37 °C for 30 min in darkness with constant shaking. Protein precipitation was performed by the addition of ice-cold acetone with vigorous vortexing, and the mixture was incubated at −20 °C for 3 h. Precipitated protein was pelleted by centrifugation at 18,000× g, 4°C for 30 min, and was gently washed with 500 μL methanol. After removing the supernatant, the protein pellet was left to air dry for 1 min, and 48 μL 50 mM Tris-formic acid (FA), pH 8.4 was added to wet the pellet. A total volume of 12 μL trypsin (Sigma-Aldrich, St. Louis, MO, USA) dissolved in 50 mM Tris-FA (0.25 μg/μL) was added to each sample to reach a final enzyme-to-substrate ratio of 1:20 (w/w), and samples were incubated in a covered thermomixer at 37 ◦C for 6 h with constant shaking. Tryptic digestion was terminated by the addition of 0.6 μL FA, and samples were centrifuged at 18,000× g, 4 °C for 30 min. The supernatant was carefully transferred to LC vials for analysis.

The LC-MS system consists of a Dionex UltiMate 3000 nano-LC system, a Dionex UltiMate 3000 micro-LC system with a WPS-3000 autosampler, and an Orbitrap Astral mass spectrometer (ThermoFisher Scientific, San Jose, CA, USA). A trapping column (100 μm ID × 360 μm OD; CoAnn Technologies, Richland, WA, USA) setting was implemented before the analytical column (75 μm ID × 30 cm L × 365 μm OD, CoAnn Technologies, Richland, WA, USA) for sample loading, cleanup and delivery. For each sample, a peptide equivalent to 500 ng protein was injected for LC-MS analysis. Detailed LC settings and information can be found in our previous publications (46). Mass spectrometry operated in data-independent acquisition (DIA) mode. MS1 spectra were acquired in the m/z range of 380 to 980 at a resolution of 180k. The maximum injection time for MS1 was set to 5 ms. Precursor ions were isolated using a 4-Th wide window and fragmented by higher-energy collision-induced dissociation (HCD) at a normalized collision energy of 25%. MS2 spectra were acquired in the Astral (AST) mass analyzer. The normalized Automatic Gain Control (AGC) target was set at 500%, with a maximum injection time of 7 ms.

For LC-MS Data Processing, the MS raw files were searched against Uniprot-SwissProt Human database (accessed December 2023) using DIA-NN (v1.8.1)1 in library-free mode. Protease was set to trypsin, the static modification was set to the carbamidomethylation of cysteine, and the dynamic modification included oxidation of methionine and acetylation of the N-terminal. Default settings were used for all other parameters.

### Seahorse XF Palmitate Oxidation Stress Test

IPSC-RPE were seeded on Matrigel-coated 96-well Seahorse plates for at least two weeks before CSE treatment. Cells were treated with CSE for one week. At 48 hours after the final CSE treatment, the cells were fed with BSA-conjugated palmitic acid (BSA-16:0) complex (102720-100, Agilent, Santa Clara, CA) diluted in XF-DMEM with glucose and L-carnitine overnight. An hour before the experiment was performed, the plates were moved to a non-CO2 incubator set at 37°C, after which the Seahorse XF Palmitate Oxidation Stress Test was performed using an XFe96 Extracellular Flux Analyzer (Agilent, Santa Clara, CA) following the manufacturer’s protocol. After the assay, cells were stained with Hoechst 33342, imaged and counted using a Cytation 1 (BioTek, Agilent, Santa Clara, CA). The oxygen consumption rate data were normalized to cell counts (per 1000 cells) and analyzed using Wave software.

### Western immunoblotting

Cells were lysed and homogenized in RIPA Buffer plus protease and phosphatase inhibitors. Protein quantification was determined using a Bicinchoninic acid protein assay. Protein (10 <g) was resolved on a 4-15% gradient polyacrylamide gel for 1 hour at 200 volts. The protein content of the gel was transferred onto a nitrocellulose membrane using the Bio-Rad Trans-Blot Turbo System. Membranes were blocked with 5% skim milk in TBST for 1 h at room temperature, then incubated overnight with primary antibodies: Perilipin-2 (Cat. #95109, Cell Signaling Technology, Danvers, MA), ABHD5 (Cat. #12201-1-AP, ProteinTech, Rosemont, IL). Horseradish peroxidase-conjugated secondary antibodies followed by Pierce ECL Western Blotting Substrate (Thermo Scientific, Waltham, MA) were used to detect proteins. Densitometric analysis was performed using Image Lab Software (Bio-Rad, Hercules, CA). Band intensities were normalized to protein load on ponceau stained membranes.

### Statistics and bioinformatic analysis

For the proteomic and lipidomics datasets, repeated measures two-way ANOVA was performed on raw values (lipidomics) or log2 transformed values (proteomics) using ezANOVA in R to analyze the main effects of CSE treatment, CFH genotype, and their interaction. Fisher’s LSD test was performed as a post-hoc analysis for pairwise comparisons. Mann Whitney U test was also used for independent analysis involving only two groups. PCA analysis was performed on Python on the selected feature set with raw values (lipidomics) or log2 transformed values (proteomics) to assess whether the experimental groups segregated. Heatmaps were generated using *pheatmap* function in R. Z-score normalization was applied to the log2-transformed expression values for visualization. Protein lists used for heatmaps were either downloaded from Gene Set Enrichment Analysis (GSEA) website for specific pathway analysis or curated based on their biological functions (GeneCards or MitoCarta 3.0). For Western blotting quantification, repeated measures two-way ANOVA were performed on the normalized expression levels (target protein/loading control). Bar graphs generated by GraphPad showed the fold change of each group relative to the control group. The significance labeled in the bar graphs were generated using normalized expression levels. Biorender (Toronto, ON) was used to create illustrations demonstrating pathways and molecular mechanisms.

## Author’s contributions

DF, JRD, and PS designed this study; PS, JH, ZG, HA, EA, and NW conducted experiments; JH, XZ, MM and JQ acquired data; PS, JH, EH and SM analyzed data; BW provided research resources; PS, DF and JH wrote the manuscript; JQ, MPA, ZG and JRD discussed data and edited the manuscript.

## Funding Support

National Eye Institute EY028554 (DAF, JRD); MPA (EY030513, American Macular Degeneration Foundation, and Research to Prevent Blindness Unrestricted Grant to Dean McGee Eye Institute).

## Acknowledgments

We thank Dr. Kevin Williams at the UCLA lipidomics core for providing technique assistance.

## Notes

### Competing Interest Statement

The authors have declared no competing interest.

