## Supplementary figures and legends for "Complement Factor H (Y402H) polymorphism for age-related macular degeneration alters retinal lipids"

### Complement Factor H (Y402H) risk polymorphism for age-related macular degeneration alters fatty acid metabolism in the retinal pigment epithelium

Peng Shang[1], Johnson Hoang[1], Elise Hong[1], Zhaohui Geng [2], Helena Ambrosino[1], Eric Abnoosian[1], Xiaoyu Zhu[3], Min Ma[3], Nova A. Wei-Navarro[4], Beau Webber [5], Jun Qu[3], Sandra R. Montezuma[6], Martin-Paul Agbaga[7], James R Dutton[2], Deborah A. Ferrington[1,8]

1. Doheny Eye Institute, Pasadena, CA 91103, USA.

2. Stem Cell Institute, University of Minnesota, Minneapolis, MN 55455, USA

3. Department of Pharmaceutical Sciences, University at Buffalo, Buffalo, NY 14214, USA

4. California Institute of Technology, Pasadena, CA 91125, USA

5. Department of Pediatrics, University of Minnesota, Minneapolis, MN 55455, USA.

6. Department of Ophthalmology and Visual Neurosciences, University of Minnesota, Minneapolis, MN 55455, USA.

7. Department of Cell Biology and Department of Ophthalmology, Dean McGee Eye Institute, University of Oklahoma Health Sciences Center, Oklahoma City, OK 73104

8. Department of Ophthalmology, David Geffen School of Medicine, UCLA, Los Angeles, CA 90095, USA

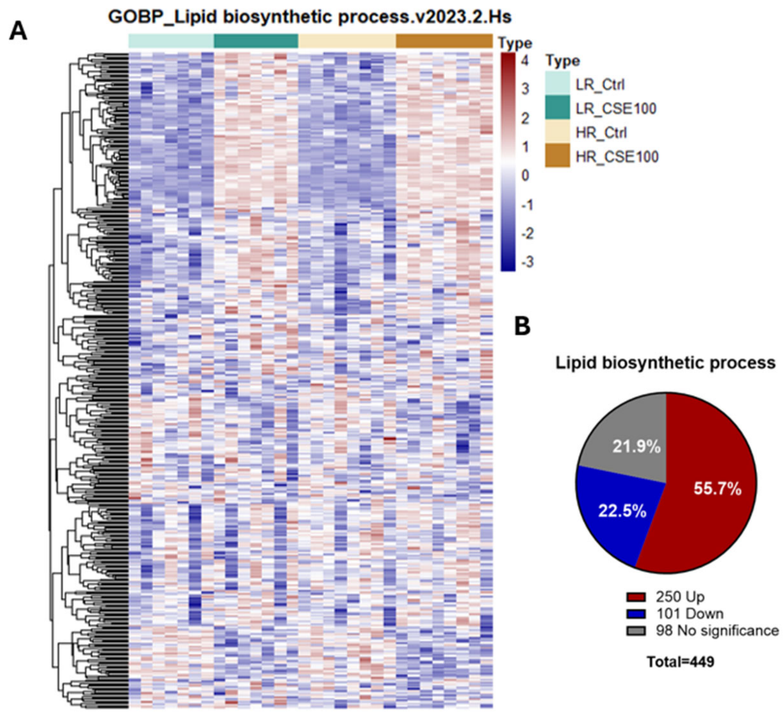


**Supplement Figure 1: CSE treatment enhances lipid biosynthesis**

(A) Heatmap with hierarchical clustering showing 449 proteins in our dataset involved in lipid biosynthetic process (gene list from GOBP_LIPID_BIOSYNTHETIC_PROCESS.v2023.2.Hs). (B) Pie chart showing the distribution of the 351 proteins that were altered significantly by CSE; 250 and 101 proteins were up-regulated or down-regulated, respectively, in both HR and LR cells. No significant change was observed in 98 proteins.

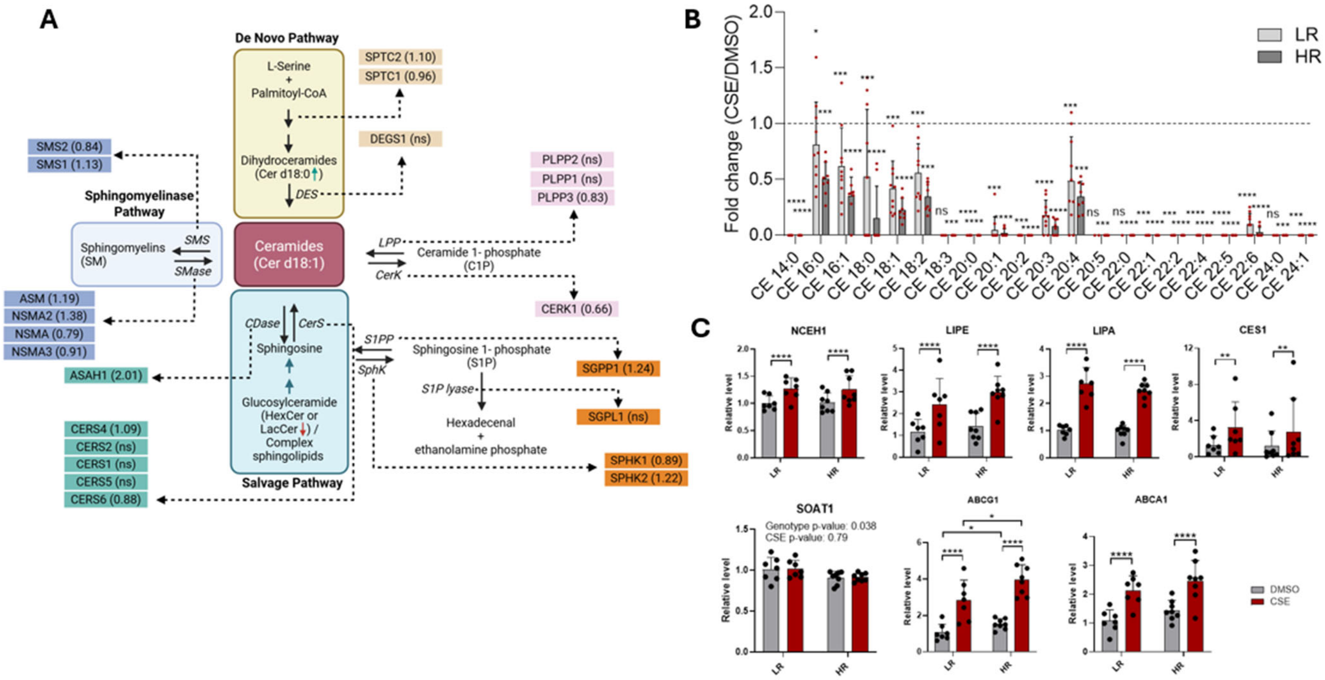


**Supplement Figure 2. CSE treatment alters ceramide and cholesterol metabolism**.

(A) Diagram of pathways involved in ceramide synthesis and metabolism (adapted from (47)). Significant changes in content of key enzymes involved in ceramide metabolism in response to chronic CSE treatment are indicated in parentheses as fold change (CSE vs. DMSO control; average of LR and HR responses). Significance was determined from the proteomics analysis using RM two-way ANOVA. (B) Bar graph showing fold change of CE species significantly altered by CSE in LR and HR cells. Analysis of each lipid species was done using RM two-way ANOVA and Fisher's LSD test. (C) Enzymes for CE hydrolysis and cholesterol transporters were significantly upregulated in both LR and HR RPE cells following CSE exposure, suggesting enhanced CE hydrolysis and cholesterol transport induced by CSE. Analysis of proteomics data was done using RM two-way ANOVA and Fisher's LSD test. Data = mean ± SD. *p<0.05, **p< 0.01, ***p<0.001, ****p<0.0001. ns = no significance.

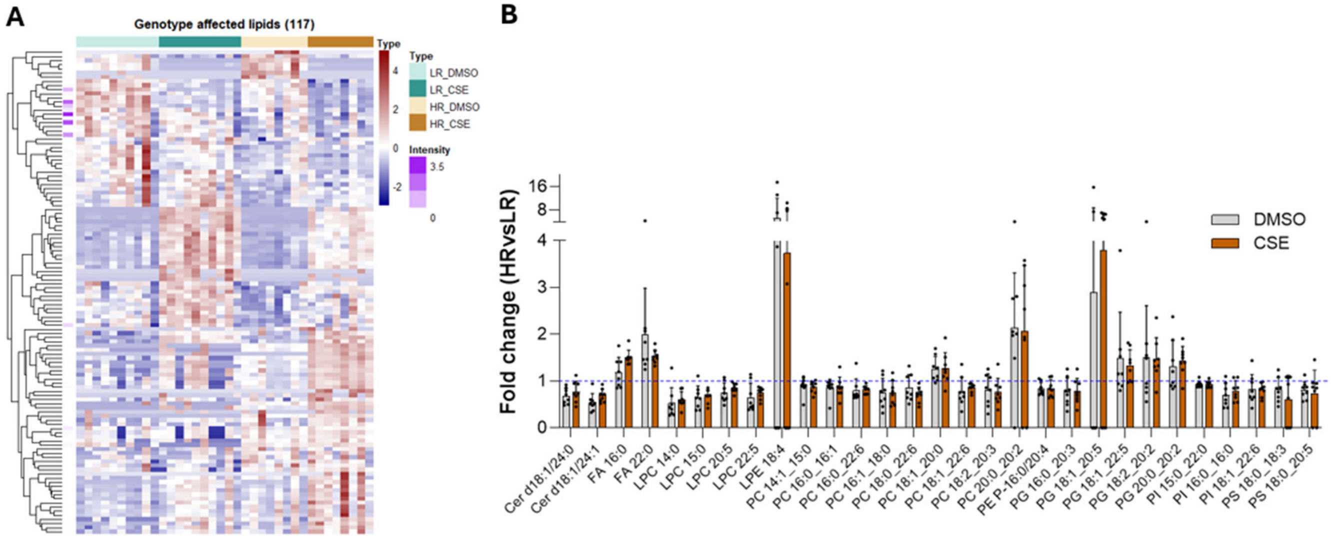


**Supplement Fig 3. Altered lipid species associated with the CFH Y402H SNP.** (A) Heatmap showing lipid species affected by CFH genotype and the interaction of CSE and genotype. (B) Fold change (HR vs LR) of Lipid species with significant genotype difference by RM two-way ANOVA, though no significant genotype differences after post hoc analysis in either basal or CSE-treated conditions.
